# *Glissandra oviformis* n. sp.: a novel predatory flagellate illuminates the character evolution within the eukaryotic clade CRuMs

**DOI:** 10.1101/2025.02.18.638917

**Authors:** Euki Yazaki, Ryo Harada, Ryu Isogai, Kohei Bamba, Ken-ichiro Ishida, Yuji Inagaki, Takashi Shiratori

## Abstract

Culturing protists offers a powerful approach to exploring eukaryotic diversity, especially for deep-branching lineages. In this study, we cultured and described a novel protist species, named *Glissan-dra oviformis* n sp., within the poorly studied and unclassified genus *Glissandra*. While an SSU rDNA gene phylogeny failed to resolve its phylogenetic placement in the eukaryotic tree, a phylogenomic analysis of 340 proteins indicated *G. oviformis* as a deep-branching member of the CRuMs clade. Prior to this study, this clade consisted of diverse heterotrophic amoeba and flagellates and lacks clear synapomorphies. Ultrastructural observations revealed that *G. oviformis* shares the characteristics with some CRuMs members, including the pellicle underlying the plasma membrane and an internal sleeve surrounding the central pair of the axoneme at the flagellar transitional region. Our findings suggest potential shared characteristics and synapomorphies for CRuMs and contribute to a deeper understanding of the character evolution within CRuMs.

## Introduction

Protists, eukaryotes excluding animals, land plants, and fungi, are a major component of the eukaryotic tree of life (eToL) and are remarkably diverse in their morphology and ecological roles (Adl. et al. 2019, Burki et al. 2020). However, compared to multicellular organisms, our understanding of unicellular protists remains limited due to their small size and difficulties in cultivation. This is well demonstrated by the fact that metabarcoding analyses hinted that diverse eukaryotic lineages, including protists, remained uncharacterized in nature (de Vargas et al. 2015, Kim et al. 2016). The limited understanding of protists suggests that there are many opportunities for new discoveries. Indeed, new protistan taxa and lineages have continued to be found, even novel supergroup-level lineages (Eglit et al. 2024, Tikhonenkov et al. 2022). Such discoveries are often based on identifying entirely novel protists or reinvestigating previously described protists with uncertain phylogenetic affiliations (here termed PUPA). These PUPA were described but not assigned to any major eukaryotic lineage due to limited morphological or molecular data. Some PUPA have been placed within well-known supergroups upon reinvestigation (Hoppenrath and Leander 2006, Shiratori and Ishida 2024, Shiratori et al. 2017), while others are revealed as distinct deep-branching lineages within the eToL (Lax et al. 2018, Yazaki et al. 2020). The long history of protist taxonomy has accumulated a substantial list of PUPA in need of reinvestigation (Adl. et al. 2019, Patterson and Zölffel 1991). Their rediscovery and characterization are essential to understanding the diversity and evolution of eukaryotes precisely. Recent phylogenetic analyses based on multiple proteins (phylogenomic analyses) have clarified the phylogenetic homes for several PUPA, such as *Microheliella*, Picozoa, and *Telonema* (Schön et al. 2021, Strassert et al. 2019, Yazaki et al. 2022). Intriguingly, morphologically dissimilar PUPA were occasionally shown to form unexpected monophyletic groups (Brown et al. 2018, Eglit et al. 2024). One prominent example is CRuMs, one of the major eukaryotic clades recently recognized (Brown et al. 2018). CRuMs was formed by three groups of PUPA: the swimming predatory flagellates (diphylleids), the non-flagellated filose amoebae (rigifilids), and the marine, tiny gliding flagellates (*Mantamonas*). Despite strong phylogenetic supports for the monophyly of CRuMs, their shared characteristics and synapomorphies remain elusive. In this study, we successfully established a culture of a new species of a previously unclassified protist genus *Glissandra*, described here as *Glissandra oviformis* n sp. A phylogenomic analysis placed *G. oviformis* as a deep-branching member of CRuMs. Furthermore, through ultrastructural observations, we inferred the shared characteristics and synapomorphies of CRuMs. These findings provide new insights into this enigmatic group and highlight the significance of PUPA in advancing our understanding of eukaryotic diversity and evolution.

## Material and methods

### Sample collection and culture establishment

A seaweed (*Halimeda* sp.) was collected in Coral Lake, the Republic of Palau (7.2510 N°, 134.3738 E°) on November 1, 2013. The seaweed was washed with IMK medium (Nihon Pharmaceutical), and the resulting wash medium was incubated at 20 °C under a 14 h light and 10 h dark cycle for five weeks. A single cell of *Glissandra oviformis* n. sp. was isolated by micropipetting and incubated with a culture of *Pedinomonas* sp. or *Bigelowiella natans* in IMK medium. The resulting two-member culture of *G. oviformis* with *Pedinomonas* sp. or B. natans was maintained at 20 °C under a 14 h light and 10 h dark cycle until April 2019.

### Light microscopy

For light microscopic observation, cells of *G. oviformis* were observed on microscope slides using a Zeiss Axio imager A2 microscope (Zeiss) equipped with an Olympus DP71 CCD camera (Olympus).

### Electron microscopy

For scanning electron microscopy, the cell suspension of *G. oviformis* was mounted on an 8.5 mm-diameter glass SEM plate (Okenshoji) treated with 0.1% (w/v) poly L-lysine. Cells sticking on the plate were prefixed with 2.5% glutaraldehyde, 0.1% osmium tetroxide (OsO4), and 0.2 M sucrose in 0.2 M sodium cacodylate buffer (CB) (pH 7.2) for 30 min and then washed with 0.2 M CB three times. The plate was placed in 0.2 M CB with 1% OsO4 for 1 h and washed with 0.2 M CB ten times. The plate was then placed in 0.2 M CB with 1% tannic acid for 30 min and washed with 0.2 M CB ten times. Finally, the plate was placed in 0.2 M CB with 1% OsO4 for 1 h again. The plate was dehydrated in a series of 15–100% (v/v) ethanol. After dehydration, the plate was placed once in a 1:1 mixture of 100% ethanol and t-butyl alcohol, twice in 100% t-butyl alcohol, and chilled in the freezer. The plate was freeze-dried using a VFD-21S (SHINKU-DEVICE) freeze drier and then mounted on aluminium stubs using carbon paste. The specimen was sputter-coated with platinum–palladium using a Hitachi E-102 sputter-coating unit (Hitachi High-Technologies) and observed using a JSM-6360F field emission SEM (JEOL). For transmission electron microscopy, the cells of *G. oviformis* were collected by centrifugation. Cell pellets were treated in a mixture of 2% (w/v) glutaraldehyde, 0.2 M sodium cacodylate, and 0.25 M sucrose for 1 h at room temperature for prefixation. Cells were then washed with 0.2 M sodium cacodylate buffer three times. Cells were then post-fixed with 1% (v/v) OsO4 in 0.2 M sodium cacodylate buffer, and dehydration was performed using a graded series of 30–100% ethanol (v/v). After dehydration, cells were placed in a 1:1 mixture of 100% ethanol and acetone, followed by pure acetone twice. Resin replacement was performed using a 1:1 mixture of acetone and Agar Low Viscosity Resin R1078 (Agar Scientific), followed by pure resin. The resin was polymerized by heating at 60°C for 12 h. Ultrathin sections were prepared on a Reichert Ultracut S ultramicrotome (Leica), double-stained with 2% (w/v) uranyl acetate and lead citrate, and observed using a Hitachi H-7650 electron microscope (Hitachi High-Technologies Corp) equipped with a Veleta TEM CCD camera (Olympus).

### DNA extraction and SSU rDNA PCR

The total DNA of the two-member culture was extracted from a cell pellet obtained by centrifuge using DNeasy plant mini kit (Qiagen Science), according to the manufacturer’s instructions. Small subunit ribosomal DNA (SSU rDNA) was amplified by PCR with forward primer 18F and reverse primer 18R (Yabuki et al. 2010). Amplifications consisted of 30 cycles of denaturation at 94 °C for 30 sec, annealing at 55 °C for 30 sec, and extension at 72 °C for 2 min. Amplified DNA fragments were purified after gel electrophoresis with a QIAquick Gel Extraction Kit (Qiagen Science) and then cloned into the p-GEM® T-easy vector (Promega). The inserted DNA fragments were completely sequenced by a 3130 Genetic Analyzer (Applied Biosystems). The SSU rDNA sequence of *G. oviformis* is deposited as xxxx in GenBank.

### SSU rDNA phylogenetic analysis

The SSU rDNA sequence of *G. oviformis* was aligned with those of 190 phylogenetically diverse eukaryotes using MAFFT v7.520 (Katoh and Standley 2013). After removing ambiguously aligned positions using BMGE v1.12 (Criscuolo and Gribaldo 2010) with default settings, 1,137 nucleotide positions remained. This alignment was subjected to the maximum-likelihood (ML) phylogenetic analysis using IQ-TREE v2.2.2.6 (Minh et al. 2020) with the SYM + I + R6 model (automatically determined by IQ-TREE’s ModelFinder) (Kalyaanamoorthy et al. 2017). ML bootstrap percentage values (MLBPs) were obtained from 100 non-parametric bootstrap replicates. Additionally, the SSU rDNA alignment was analyzed using Bayesian phylogenetic method with PHYLOBAYES-mpi v1.8a (Lartillot and Philippe 2004, 2006, Lartillot et al. 2007). In this analysis, two MCMC runs were conducted for 100,000 cycles with a burn-in of 25,000 cycles. A consensus tree with branch lengths and Bayesian posterior probabilities (BPPs) was then calculated from the remaining trees.

### RNA extraction and RNA-seq

The total RNA of the two-member culture was extracted from a cell pellet obtained by centrifuge using the Trizol reagent (Life Technologies) following the manufacturer’s instructions. The cDNA library construction and paired-end sequencing (125 bp per read) with HiSeq2000 (Illumina) were performed at Eurofins Genomics. For *G. oviformis*, 3.3 × 107 paired-end 125 bp reads (8.2 Gb in total) were obtained and then performed quality filtering using Trimmomatic v0.39 (Bolger et al. 2014) with the following parameters: LEADING:30, TRAILING:30, SLIDING-WINDOW:4:25, and MINLEN:20. The filtered reads were then mapped to the CDS of *B. natans* (Bigna1) (Curtis et al. 2012) downloaded from EnsemblProtists (https://ftp.ensemblgenomes.ebi.ac.uk/pub/protists/release-59/fasta/bigelowiella_natans/cds/) using HISAT2 v2.2.1 (Kim et al. 2019). Unmapped reads were assembled using Trinity v2.15.1 (Grabherr et al. 2011, Haas et al. 2013) to obtain unique contigs of 81,591. The contigs were translated into protein sequences using TransDecoder v5.7.1 (https://github.com/TransDecoder/TransDecoder), resulting in the identification of 376,986 protein sequences.

### Phylogenomic analyses

To elucidate the phylogenetic position of *G. oviformis*, we prepared a phylogenomic alignment by updating an existing one comprising 351 proteins (Harada et al. 2024). For each protein, we added homologous sequences retrieved by BLASTp (with an E-value cut-off of 1030) from the newly generated *G. oviformis* protein sequences (see above), as well as sequences from the provorans, anaeroamoebids, and other organisms (total 50 species including *G. oviformis* added, described in Table S1). Individual single-protein alignments were generated using MAFFT v7.520 with the L-INS-i algorithm, followed by manual correction and automated exclusion of ambiguously aligned positions using BMGE v1.12. Each single-protein alignment was subjected to a preliminary phylogenetic analysis using FASTTREE v2.1.11 (Price et al. 2010) under the LG + Γ model. The resulting approximately maximum-likelihood trees with SH-like local supports were examined to identify alignments with aberrant phylogenetic signals that strongly disagreed with well-established monophyletic groups in the eToL, including Opisthokonta, Amoebozoa, Alveolata, Stramenopiles, Rhizaria, Rhodophyta, Chloroplastida, Glaucophyta, Haptophyta, Cryptista, Jakobida, Euglenozoa, Heterolobosea, Diplomonadida, Parabasalia, and Malawimonadida. Eleven out of the 351 single-protein alignments were found to violate these criteria and were excluded from further phylogenomic analyses. The remaining 340 single-protein alignments (see Table S1) were concatenated into a single phylogenomic alignment comprising 132 taxa with 88,397 unambiguously aligned amino acid positions. The coverage for each single-protein alignment is summarized in Table S1. The final alignment comprising 340 proteins from 132 taxa (340-protein alignment) was subjected to the ML analysis using IQ-TREE v2.2.2.6 with the LG + Γ + F + C60 model. The robustness of the ML phylogenetic tree was assessed through ultra-fast ML bootstrap approximation with the LG + Γ + F + C60 model (1,000 replicates) and a non-parametric ML bootstrap analysis with the LG + Γ + F + C60 + PMSF (posterior mean site frequencies) model (200 replicates) (Wang et al. 2018). The ML tree inferred with the LG + Γ + F + C60 model served as the guide tree for the non-parametric bootstrap analysis incorporating PMSF. Additionally, Bayesian phylogenetic analysis was conducted using the CAT + GTR model with PHYLOBAYES-mpi v1.8a. In this analysis, two MCMC runs were conducted for 5000 cycles. After discarding the trees from the first 1250 cycles as burn-in, the consensus tree, including branch lengths and Bayesian posterior probabilities (BPPs), was calculated from the remaining trees. In addition, we evaluated each bipartition using AUTOEB v1.0.0 (Bamba et al. 2024), in which the two alternative trees on each bipartition were generated by nearest-neighbor-interchange followed by an approximately unbiased (AU) test comparing the original (ML) tree and the two alternative trees. The settings of AUTOEB were set as the default, and the substitution model was the same as the ML phylogenetic analysis of the 340-proteins alignment. We evaluated the impact of fast-evolving positions within the 340-protein alignment. Substitution rates for each position were calculated on the ML tree by IQ-TREE v2.2.2.6, and four sets of subalignments were created from the original alignment by sequentially removing the top 20%, 40%, 60%, and 80% fastest-evolving positions. The original alignment and four sub-alignments were individually subjected to the ultra-fast bootstrapping approximation under the LG + Γ + F + C20 model using IQ-TREE v 2.2.2.6.

### Environmental SSU rDNA sequences

To examine the abundance and distribution of *G. oviformis*, we searched for sequences similar to its SSU rDNA sequence in the EukBank global dataset (Berney et al. 2023) that contains V4 sequences in SSU rDNA from 12,672 samples collected across marine and terrestrial environments. A BLASTn search was conducted using the *G. oviformis* SSU rDNA sequence as a query, and hits with over 90% length of the subject sequence were picked.

## Results

### Morphology and ultrastructure of *Glissandra oviformis* n. sp

Cells of *Glissandra oviformis* n. sp. were oval or ovoid, measuring approximately 4.8 (3.7–6.5) µm in length and 3.5 (2.4–5.3) µm in width (n = 36) (figure 1). Each cell possessed two flagella of subequal length, 2.5 to 4 times the cell body length (figure 1a, c, d). These flagella are inserted longitudinally from a depression near the anterior end on the ventral side of the cell (figure 1a, c, d). Cells exhibited gliding motility, with most of both flagella attached to the substrate. During gliding, the anterior flagellum extends straight forward, while its tip detaches from the substrate and moves back and forth. The posterior flagellum extended posteriorly, attaching to the substrate along its entire length. Cells possessed a distinct aperture at the posterior side of the flagellar depression. Some cells contained prey-derived green or orange food vacuoles (figure 1e, f). In the scanning electron microscopy, the aperture and the flagellar depression continued, and their rims were swollen except for the left edge of the flagellar depression (figure 2a, b). No appendages, such as mastigonemes or scales, were observed on either the flagella or the cell surface (figure 2a, b). Transmission electron microscopy showed *G. oviformis* has a nucleus at the posterior region of the cell (figure 2c). The nucleus contained two nucleolus-like structures with different electron densities (figure 2d). Virus-like particles covered by envelope were occasionally observed in the nucleus and cytoplasm (figure 2d, S1a). Cells have several mitochondrial profiles with lamellar cristae (figure 2c, e). An inflated Golgi apparatus was situated near the base of the basal bodies (figure 2c). In the Golgi cisternae, bridges that link the two closely opposed membranes were observed (figure S1b). These bridges appear to be placed on the same plane (figure S1c). A spherical microbody was observed in the cell (figure 2e). Lipid droplets were occasionally observed in the cytoplasm (figure S1d–g). A single-layered pellicle underlying the plasma membrane, except in the region of the flagellar depression and the aperture, was observed (figure 2f). The pellicle exhibited inward curling at the rim of the aperture and the right edge of the flagellar depression (figure 2g, h). The basal bodies were arranged at an acute angle of approximately 25–40°(figure 2i). Both anterior and posterior basal bodies possess dorsally directed fibers (figure S1d–g). A spiral fiber with 4–6 gyres was present in the flagellar transitional region (figure 2j, l, m). In the spiral fiber, a dense plate-shaped axosome was sandwiched by less dense amorphous materials (figure 2j). An internal sleeve surrounded the central pair of the axoneme just above the spiral fiber (figure 2j–l). The basal bodies had three microtubular roots. Following the terminology of Yubuki and Leander (2013), each microtubular root was labeled R1, R2, and R3. R1 emerged from the anterior side of the posterior basal body (figure 2i). It comprised five microtubules and an electron-dense material associated with its proximal end (figure 2i, n). R1 initially directed leftward but then curved posteriorly, terminating near the anterior edge of the aperture (figure 2o, S2). R2 consisted of four microtubules arranged in a U-shape and emerged from the posterior side of the posterior basal body (figure 2n, S2c–h, S3g, h). R2 directed toward the anterior edge of the aperture and further extended to line its left edge (figure 2c, o, S2). R3 emerged from the anterior side of the anterior basal body, composed of three microtubules (figure 2c, o, S2a, S3a). R3 directed dorsally, passing just beneath the anterior cell surface, and lacked secondary microtubules (figure 2c, o).

**Figure 1.**
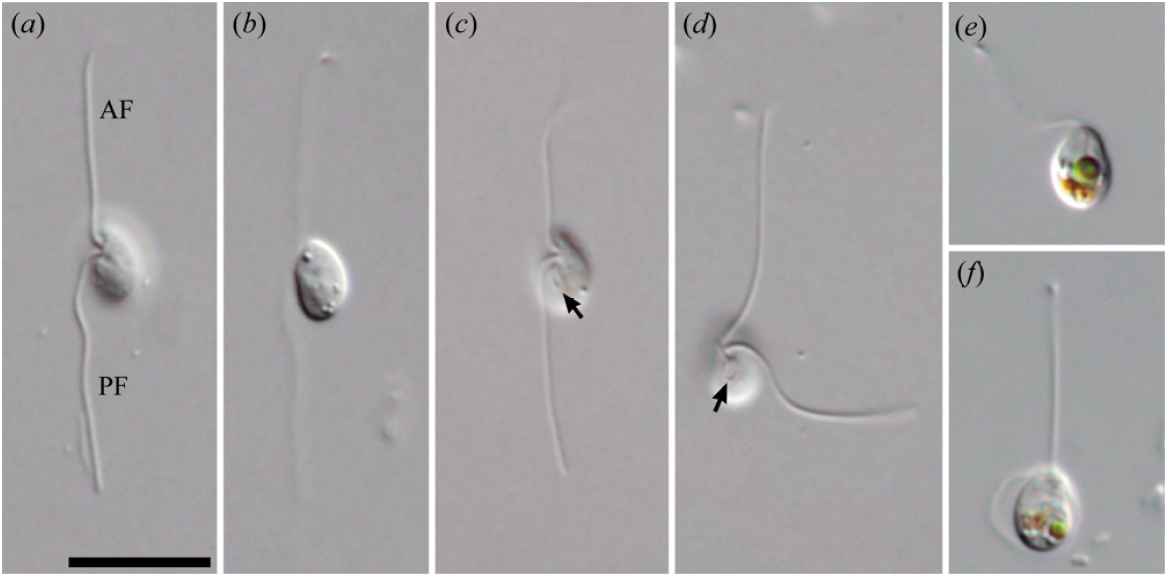
Differential interference contrast micrographs of *Glissandra oviformis* n. sp. (**a, b**) Gliding cells. (**c, d**) Gliding cells showing ventral apertures. (**e, f**) Gliding cells showing food vacuoles. AF, anterior flagellum; PF, posterior flagellum. Arrows indicate ventral apertures. Scale bar = 10 µm.

**Figure 2.**
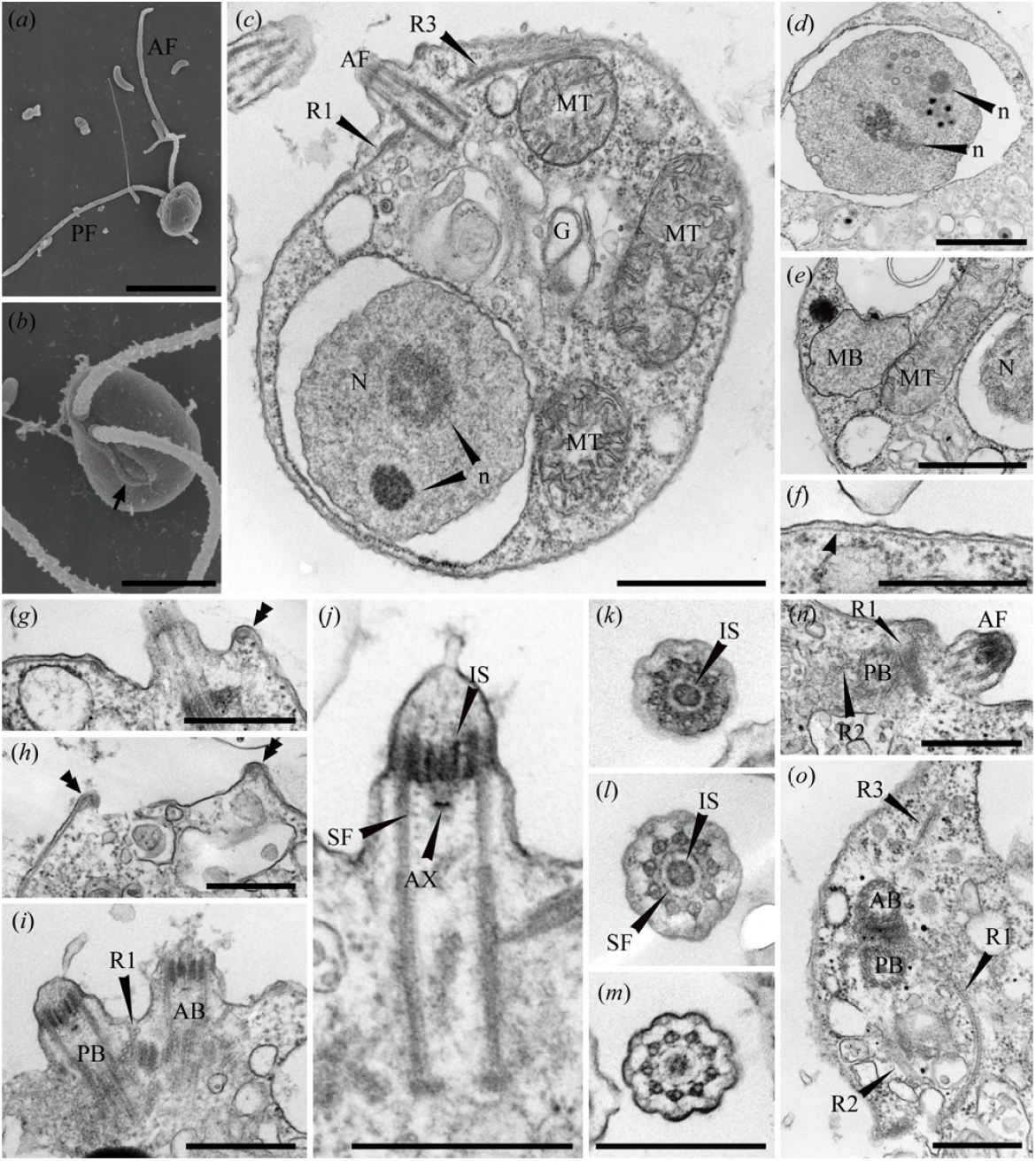
Scanning and transmission electron micrographs of *Glissandra oviformis* n. sp. (**a**) Scanning electron micrograph (SEM) of a whole cell. **b** SEM of a flagellar depression and a ventral aperture. **c** Transmission electron micrograph (TEM) of the whole cell **d** TEM of a nucleus and virus-like particles in the nucleus. **e** TEM of a microbody and a mitochondrion. **f** TEM of a pellicle underlying the plasma membrane. **g** TEM of inward curling of pellicle at the right edge of the flagellar depression. **h** TEM of inward curling of pellicle at the rim of a ventral aperture. **i** TEM of the longitudinal section of two basal bodies. **j** TEM of the longitudinal section of anterior basal body. **k** TEM of a transverse section of a flagellar transitional region at the level of an internal sleeve. **l** TEM of a transverse section of a flagellar transitional region at the level between the internal sleeve and a spiral fiber. **m** TEM of transverse section of a flagellar transitional region at the level of the spiral fiber. **n** TEM of the proximal end of the R2. **o** TEM of a transverse section of two basal bodies. AB, anterior basal body; AF, anterior flagellum; AX, axosome; G, Golgi apparatus; IS, internal sleeve; MB, microbody; MT, mitochondrion; N, nucleus; n, nucleolus-like structure; PB, posterior basal body; PF, posterior flagellum. SF, spiral fiber. An arrow indicates ventral aperture. An arrowhead indicates pellicle. Double arrowheads indicate rims of ventral aperture or flagellar depression. Scale bars: (a) = 5µm, (b–e) = 1 µm, (f–h, j, l–o) = 500 nm.

### Phylogenomic analyses nominated *G. oviformis* as a member of CRuMs

We gathered 190 phylogenetically diverse eukaryotic SSU rDNA sequences and conducted the phylogenetic analysis using SSU rDNA to determine the phylogenetic position of *G. oviformis*. However, the phylogenetic placement of *G. oviformis* remained unresolved in the SSU rDNA phylogenetic tree. Specifically, *G. oviformis* branched as a sister lineage to Cyanidiaceae, though this bipartition lacked statistical support (an MLBP of 18%, figure S4). To further resolve the phylogenetic position of *G. oviformis*, we performed phylogenomic analyses using a 340-protein alignment from 132 taxa representing all the major eukaryotic groups. The resulting phylogenomic tree robustly supported the monophyly of all the major eukaryotic groups (figure 3a). In these analyses, *G. oviformis* was placed within the CRuMs, which includes *Mantamonas plastica, Collodictyon triciliatum, Diphylleia* sp., and *Rigifila ramosa*. The assemblage comprising the previously known members of CRuMs and *G. oviformis* received full statistical support (figure 3a). Within this clade, *G. oviformis* branches after *M. plastica*, the deepest branching taxon in the CRuMs, followed by *Rigifila* and two diphylleids. Each bipartition within this assemblage in the ML tree also received full statistical support and remained consistently supported even after the removal of fast-evolving positions (figure 3b). Thus, we conclude that *G. oviformis* is a member of CRuMs and represents a previously overlooked subclade in CRuMs. We here referred to the clade consisting of glissandrids, rigifilids, and diphylleids as “GRiDi.” It is worth noting the incongruity between the ML and Bayesian analyses. The CRuMs clade enclosing *G. oviformis* is positioned as the sister group to Amorphea (this assemblage is known as Podiata; Cavalier-Smith 2013), with this relationship being robustly supported in the assessments using the ML method (figure 3a). This relationship remains consistent even after excluding fast-evolving positions (figure 3b). In contrast, Bayesian phylogenetic analyses placed the clade of parabasalids and aneroamoebas as the sister to Amorphea, thereby failing to recover the Podiata (figure S5). The bipartition uniting Amorphea, parabasalids, and aneroamoebas together received an inconclusive BPP of 0.65, suggesting that the precise relationship among Amorphea, CRuMs, and the clade of parabasalids and aneroamoebas remained uncertain in Bayesian analysis.

**Figure 3.**
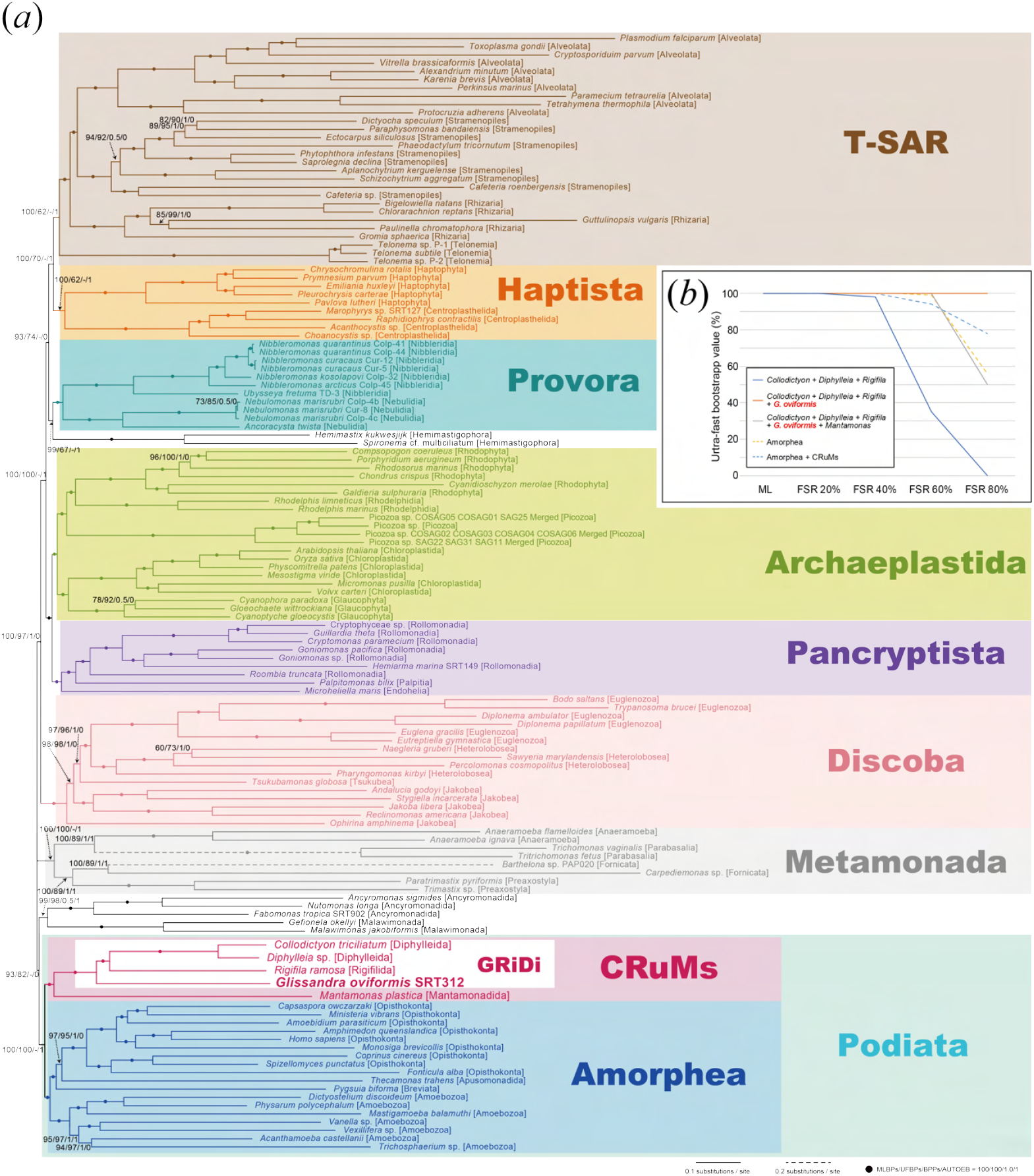
Phylogenomic analyses of a 340-protein alignment. The ML tree was inferred from a 340-protein alignment (comprising 132 taxa and 88,397 amino acid positions) as shown in (**a**). The same alignment was subjected to Bayesian analysis and the resultant BPPs were mapped on the ML tree. The detailed Bayesian tree is provided in figure S4. For each node, MLBPs, ultrafast bootstrap values (UFBPs), Bayesian posterior probabilities (BPPs), and results from AUTOEB analysis are presented. Nodes marked with dots indicate MLBPs = 100%, UFBPs = 100%, BPPs = 1.0, and the calls from AUTOEB as “resolved” (marked by “1”). The results from the analyses of the 340-genes alignment processed by fast-evolving position removal (FPR) are presented in (**b**). Ultrafast bootstrap analyses were performed on the original 340-genes alignment and alignments after removing the top 20%, 40%, 60%, and 80% of fastest-evolving positions using IQ-TREE 2.2.2.6 under the LG + Γ + F + C20 model. Solid lines in blue, orange, and gray indicate UFBPs for the monophyly of (*Collodictyon* + *Diphylleia* + *Rigifila*), (*Collodictyon* + *Diphylleia* + *Rigifila* + *G. oviformis*), and (*Collodictyon* + *Diphylleia* + *Rigifila* + *G. oviformis* + *Mantamonas*), respectively. Dashed lines in yellow and light blue indicate UFBPs for the monophyly of Amorphea, Amorphea + CRuMs, respectively.

### Putative marine habitat of *G. oviformis* and its relatives

A BLASTn search against the Eukbank dataset identified 13 amplicon sequence variants (ASV) aligned to the SSU rDNA of *G. oviformis* in over 90% of their length. One of the 13 ASVs matched perfectly with the SSU rDNA of *G. oviformis* (table S2). This ASV was assigned the rhodophyta “*Porphyra capensis*” in the Eukbank dataset but the sequence similarity between the ASV and the SSU rDNA of *P. capensis* deposited in the GenBank database (AY766361) was only 74.4%. Therefore, this ASV originated from *G. oviformis*, not the rhodophyte. We could not conclude the precise origins of the other 12 ASVs, as they did not match the *G. oviformis* sequence or any of the rDNA sequences in the GenBank with high identity (table S2). The *G. oviformis* ASV was not detected in any terrestrial sample but in 47 samples collected from marine sediment and water columns with a depth range from the surface to 800 m and distributed across the Mediterranean Sea, Indian Ocean, Pacific Ocean, and Atlantic Ocean. The *G. oviformis* ASV consists of 5,679 reads; most of them (5,603 reads) originated from a seawater sample collected from the Indian Ocean and reached over 1% of the total reads. In the other 46 samples, the reads derived from *G. oviformis* occupied less than 0.03% of the total reads (table S3). This low abundance of *G. oviformis* ASV reads can be interpreted as this species being scarce in marine environments on top of its ecological role as a predator.

## Discussion

### Glissandra oviformis n. sp. as a new species of the genus Glissandra

In our light microscopic observation, *Glissandra oviformis* n. sp. showed characteristics consistent with other *Glissandra* species, gliding flagellates possessing anteriorly and posteriorly directed flagella and the back-and-forth movement of the anterior flagellar tip. *Glissandra* contains two species: *G. innuerende*, the type species, and *G. similis* (Lee 2006, Patterson and Simpson 1996). *G. innuerende* resembles *G. oviformis* in attaching both flagella to the substrate and possessing an aperture (as depicted in Fig 8b of Patterson and Simpson 1996) on the ventral side of the cell. However, in *G. innuerende*, the two flagella were inserted laterally, whereas those of *G. oviformis* were inserted longitudinally. The longitudinal arrangement of flagella in *G. oviformis* was confirmed through our light and electron microscopy, and no variations were observed within the strain. Additionally, there is a slight difference in cell shape, with *G. innuerende* being almost spherical while *G. oviformis* is oval. Another species, *G. similis* differs from both *G. innuerende* and *G. oviformis*, as it possesses a ventral groove instead of a ventral aperture and attaches to the substrate using the tip of the posterior flagellum (Lee 2006). Because *G. oviformis* can be distinguished from the two described species of *Glissandra*, we propose it as a new species of the genus *Glissandra*.

### *G. oviformis* reveals character evolution in CRuMs

CRuMs, encompassing three lineages [Diphylleida (Collodictyonida), Rigifillida, and *Mantamonas*], has thus far been represented by only 12 species, making it a relatively small group despite the clade being compatible with a supergroup. In this study, our phylogenomic analyses have clarified the phylogenetic position of *G. oviformis* within the CRuMs clade (figure 3a, b, S5). Combined with the concrete phylogenetic placement of *G. oviformis* within CRuMs, the detailed morphological data from this organism provided the first ground for discussing the character evolution in this clade. Each of the three CRuMs lineages recognized previously exhibit distinct morphological characteristics. Diphylleida comprises freshwater swimming flagellates with a prominent ventral groove (Brugerolle 2006; Brugerolle et al. 2002). Rigifilida is a group of non-flagellated filose amoebae, including *Rigifila* and *Micronuclearia* (Mikrjukov and Mylnikov 2001; Yabuki et al. 2013). *Mantamonas* represents asymmetric, small bacterivorous gliding flagellates consisting of three species, but ultrastructural data for this group are currently unavailable (Blaz et al. 2023; Glücksman et al. 2010). Here, we compare the morphologies and ultrastructures of *G. oviformis* with those of other members of CRuMs and other potentially related taxa to infer the shared characters and synapomorphies of the group. The presence of pellicles underlying the plasma membrane is observed in the currently recognized members of CRuMs, except for *Mantamonas*, with no ultrastructural data available. Within Rigifilida, single-layered (*Micronuclearia*) or double-layered (*Rigifila*) pellicles support the plasma membrane, with the exception of an aperture through which pseudopodia emerge (Mikrjukov and Mylnikov 2001; Yabuki et al. 2013). Although the original ultrastructural studies of diphylleids did not mention a pellicle, Cavalier-Smith (2013) pointed out the presence of pellicles in *Collodictyon* and *Diphylleia* (see Figures 2e, g, h in Brugerolle et al. 2002 and Figures 24, 28 and 32 in Brugerolle and Patterson 1990). The pellicles underlying the plasma membrane appear to extend from the rim of the ventral groove to the dorsal side of the cell. However, it remains unclear whether the entire dorsal surface is covered. Similar to *Micronuclearia* and diphylleids, *G. oviformis* also possesses a single-layered pellicle just beneath the plasma membrane, with exceptions at the flagellar depression and ventral aperture. The widespread occurrence of pellicles suggests that the common ancestor of GRiDi possessed this structure (figure 4). We demand the ultrastructural information on *Mantamonas* to clarify whether the pellicle is a shared characteristic among all CRuMs lineages. Intracellular pellicles are also reported in apusomonads and ancyromonads (Heiss et al. 2011, Karpov and Zhukov 1986, Molina and Nerad 1991). Ultrastructural studies on ancyromonads reveal a single-layered pellicle underlying the plasma membrane, with exceptions at the base of the flagellar pockets and a channel extending from the posterior flagellar pocket (Heiss et al. 2011, Blaz et al. 2023). On the other hand, apusomonad pellicle structures vary across studies, ranging from thin, single-layered structures resembling that of Ancyromonas (Heiss et al., 2013) to complex, five-layered structures (Karpov and Mylnikov, 1989; Karpov and Zhukov, 1986). These pellicles underline the dorsal side of the cell. Apusomonads and ancyromonads were previously classified together in phylum Apusozoa along with rigifilids based on the presence of pellicles. (Cavalier-Smith et al. 2008, Yabuki et al. 2013). However, recent phylogenomics placed apusomonads within Obazoa, along with Opisthokonta and Breviatea (Brown et al. 2013, Brown et al. 2018). Ancyromonads, on the other hand, remain unclassified but occasionally show a close relationship to Podiata, together with Malawimonadida (Brown et al. 2013; Brown et al. 2018, Harada et al. 2024, Torruella et al. 2024, Strassert et al. 2019). Our ML phylogenomic tree also suggested that ancyromonads (along with malawimonads) are related to Podiata. However, this topology was not supported by the AU test conducted through AUTOEB and fast-evolving position removal analyses (figures 3a b), nor was it reconstructed using the Bayesian method (figures 3a, b, S5). These morphological features and molecular phylogenies may suggest that pellicle does not originate within CRuMs but is a more ancestral character. Structures in flagellar transitional regions and fibrillar or microtubular roots associated with the basal bodies have been used as significant characteristics for higher classification of protists (Yubuki and Leander 2013; Yubuki et al. 2016). While rigifilids are amoebae that lack flagella, and data on the ultrastructure of *Mantamonas* absent, the flagellar apparatus of diphylleids has been extensively studied (Brugerolle et al. 2002, Brugerolle 2006, Brugerolle and Patterson 1990). In all genera of diphylleids, a sleeve surrounding the central pair of the axoneme has been reported within the flagellar transitional region (Brugerolle 2006). Because we observed the sleeve in *G. oviformis* as well, this feature is the clear synapomorphy for GRiDi or CRuMs (figure 4). The flagellar apparatus of diphylleids shares similarities with that of *G. oviformis* in terms of the microtubular roots, which consist exclusively of R1, R2, and R3 (corresponding to lvF, rvF, and mL in Brugerolle and Patterson 1990 respectively). Both R1 and R2 are directed posteriorly, and R3 is directed dorsally through a position just beneath the anterior surface of the cell. The absence of both the singlet root and R4, which are widely distributed within eukaryotes (Yubuki et al. 2013), could potentially serve as a taxonomic trait for CRuMs. However, the flagellar apparatus of diphylleids is more complex than that of *G. oviformis* in other aspects. Each microtubular root in diphylleids has an associated electron-dense structure, and R3 possesses well-developed secondary microtubules referred to as a dorsal fan. Additionally, microfibrillar fans are associated with each basal body (Brugerolle et al. 2002, Brugerolle 2006, Brugerolle and Patterson 1990). The differences in flagellar apparatus between *G. oviformis* and diphylleids may be related to differences in cell size, mode of movement, and feeding behavior between these two groups. The shapes of mitochondrial cristae vary among CRuMs. Diphylleids possess tubular cristae, while rigifilids have lamellate (M. podoventralis) or irregularly flat (R. ramose) cristae (Brugerolle 2006; Brugerolle et al. 2002; Mikrjukov and Mylnikov 2001; Yabuki et al. 2013). *G. oviformis* has lamellar cristae similar to those of R. ramosa. This diversity suggests that cristae shape is not a conserved feature within CRuMs. Yabuki et al. (2013) proposed a potential relationship between rigifilids and diphylleids based on the presence of pseudopodia. Since *Mantamonas* also possesses pseudopodia, this feature was likely present in the common ancestor of CRuMs (Glücksman et al. 2010). We did not observe pseudopodia in *G. oviformis*. Therefore, pseudopodia may have been lost in this species or might only appear during specific life stages, such as predation.

**Figure 4.**
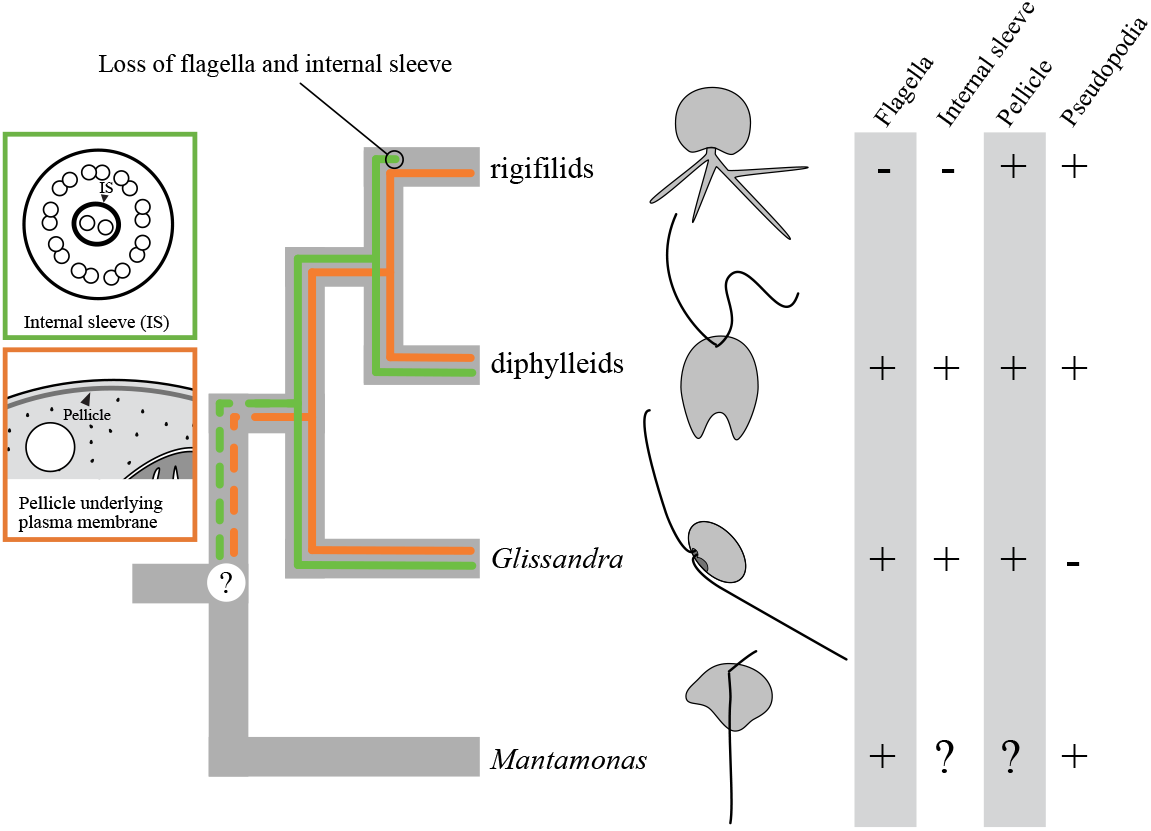
The putative character evolution within CRuMs based on the tree of CRuMs inferred from a 340-protein alignment. Green and orange lines represent the presence of the internal sleeve and the pellicle, respectively.

### Perspectives toward understanding the diversity of CRuMs

It is too naive to assume that the genuine diversity of CRuMs is sufficiently represented by Diphylleida (Collodictyonida), Rigifillida, *Mantamonas*, and *Glissandra*. We anticipate that (potentially a number of) as-yet-undescribed members of this clade may remain to be found in natural environments. Despite the diversity of CRuMs being a significant piece of information to picture the eToL, the sequencing analyses based on environmental DNA (eDNA) or cDNAs/genomes amplified from the cells or a single cell isolated from environmental samples are unlikely practical tools to identify novel CRuMs members in the future. Recent advances in metabarcoding have enabled the identification of novel lineages without the extensive time and effort required for establishing laboratory cultures. Marine Stramenopiles (MAST) and Marine Alveolata (MALV) are prime examples, significantly contributing to the understanding of the diversity within Stramenopiles and Alveolata lineages (Massana et al. 2004; López-García et al. 2001). In particular, the intra-lineage diversity of organisms that are difficult to cultivate, such as Picozoa, has been substantially expanded through metabarcoding efforts (Huber et al. 2024, Kim et al. 2016, Not et al. 2007). Meanwhile metabarcoding amplifies hyper-variable regions approximately 200–400 bp in length (e.g., the V4 or V9 regions of SSU rDNA) from the organisms present in the environment sample of interest. When the nucleotide sequence of an amplicon is identical (or very similar) to the SSU rDNA sequence linked to the previously known organism with a cellular identity, we can conclude that the environmental sample examined contained the particular organism. In other words, due to the limited phylogenetic information in the short amplicons, those bearing no similarity to any known SSU rDNA sequences provide little information on their cellular origins. Moreover, there are potentially a large number of organisms that have difficulty in amplifying the regions to be targeted in typical surveys based on eDNA samples, and such organisms may remain undetected (e.g., Euglenozoa and Metamonada, Novák et al. 2024). The above-mentioned issues in the experiments based on short amplicons can be overcome partially by amplifying, sequencing, and phylogenetically analyzing long amplicons (approximately 1 kb), including both variable and conserved regions (Tedersoo et al. 2017; Latz et al. 2022). However, as demonstrated in figure S4, the currently recognized members of CRuMs did not form a single clade in the phylogenetic analysis using (nearly) full-length SSU rDNA sequences. Even closely related species, such as *Diphylleia* and *Collodictyon*, form distinct clades in the SSU rRNA phylogeny. Thus, phylogenetic analyses of full-length SSU rDNA sequences are most likely insufficient to identify novel members of CRuMs. Only phylogenomic analyses based on high-quality transcriptome data unveiled novel members of CRuMs and their internal relationships with confidence. We can now obtain transcriptome and genome data from a small number of cells (even a single cell) isolated from environment samples, and such large-scale sequence data have been regarded as an efficient method for resolving various subtrees of the eToL (e.g., Kolisko et al. 2014). However, it is difficult to tell which cell is a novel CRuMs member under a microscope, as the cell morphology varies largely among the currently known members in this clade. Thus, the survey explicitly targeting novel CRuMs members, followed by single cell-based, largescale sequencing, is likely infeasible. Unfortunately, there is no experimental shortcut for the studies exploring the diversity and evolution of CRuMs. We must keep searching for previously undescribed eukaryotes in various types of natural environments to encounter novel organisms belonging to CRuMs in the future. It is also possible that, as *Glissandra*, novel CRuMs members might be overlooked among the previously described protists with uncertain phylogenetic affiliations.

## Taxonomic Summary

### CRuMs

Order Glissandrida ord. nov. Diagnosis: Heterotrophic biflagellates. Flagellar transitional region includes the sleeve that surrounds the central pair of flagellar microtubules. Single-layered pellicle that underlying plasma membrane except for a flagellar insertion and a ventral aperture. Mitochondrial cristae with irregularly flat shape. ZooBank LSID:xxxxxxxxx

#### Family Glissandridae fam. nov. Diagnosis

Same as order. Type genus: *Glissandra* ZooBank LSID: xxxxxxxxx Genus *Glissandra* Emended diagnosis: Gliding biflagellates. Anterior flagellum directed anteriorly, and the posterior flagellum directed posteriorly in gliding cell. Only the tip of the anterior flagellum moves back and forward during gliding. Ventral aperture or ventral groove present. ZooBank LSID:xxxxxxxxx *Glissandra oviformis* n. sp. Diagnosis: Cells are oval or ovoid, 3.7–6.5 µm in length and 2.4–5.3 µm in width. Two subequal flagella of 2.5–4 times the cell length inserted longitudinally from a subapical depression at ventral side of the cell. Both flagella lie against the substrate in gliding cells. Hapantotype: One microscope slide (TNS AL-xxxxx), deposited in the herbarium of the National Museum of Nature and Science (TNS), Tsukuba, Japan.

#### Paratype

One EM block (TNS AL-xxxxxx) deposited in the TNS. These cells are derived from the same sample as the hapantotype. DNA sequence: Small subunit ribosomal RNA gene, xxxxxx. Type locality: A sample of seaweed (*Halimeda* sp.) in Coral Lake, Palau (latitude = 7.2510 N°, longitude = 134.3738 E°) Collection date: November 1, 2013.

#### Etymology

The specific epithet “oviformis” (egg-shaped) refers to the cell shape of this organism. ZooBank LSID:xxxxxxxxx

## Ethics

Permits for collecting marine resources were obtained from the Ministry of Natural Resources, Environment, and Tourism, the Republic of Palau (permit no. RE-13-15 at 2013).

## Data accessibility

Available on requests to EY.

## Acknowledgments

We thank Dr. Naoto Hanzawa at Yamagata University for trip management for the sampling in the Republic of Palau. R.H. was supported by JSPS Overseas Research Fellowships. References

## Supplementary figures

**Figure S1.**
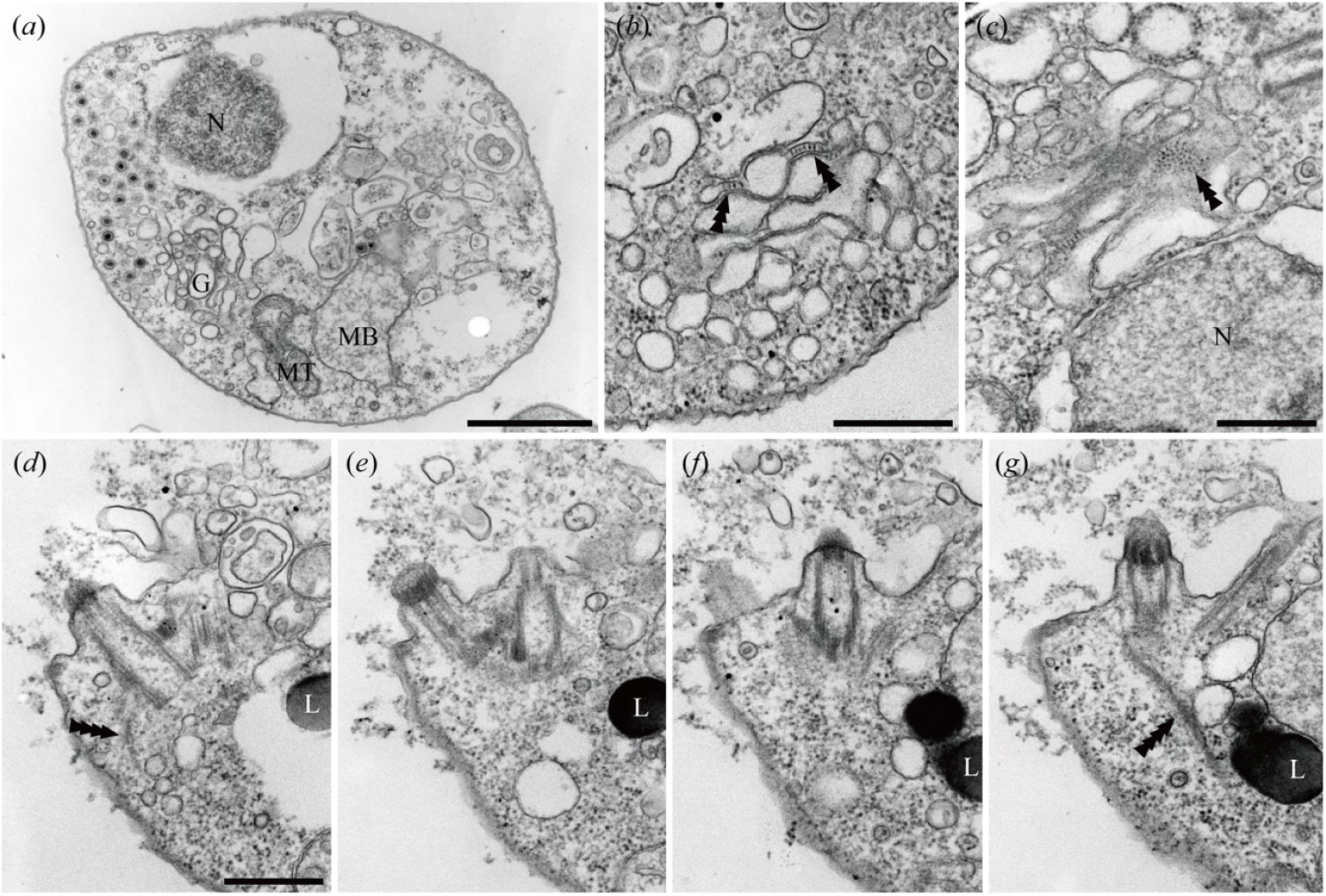
Transmission electron micrographs (TEM) of *Glissandra oviformis* n. sp. (**a**) TEM of the whole cell showing virus-like particles in cytoplasm. (**b, c**) TEM of Golgi apparatus showing bridges that link the two closely opposed membranes. (**d-g**) Selected serial TEM of fibers emerged from the basal bodies. L, lipid droplet; MB, microbody; MT, mitochondrion; N, nucleus. Triple arrowheads indicate bridges of the Golgi apparatus. Quadruple arrowheads indicate fibers emerged from the basal bodies. Scale bars: (a) = 1 µm, (b–g) = 500 nm

**Figure S2.**
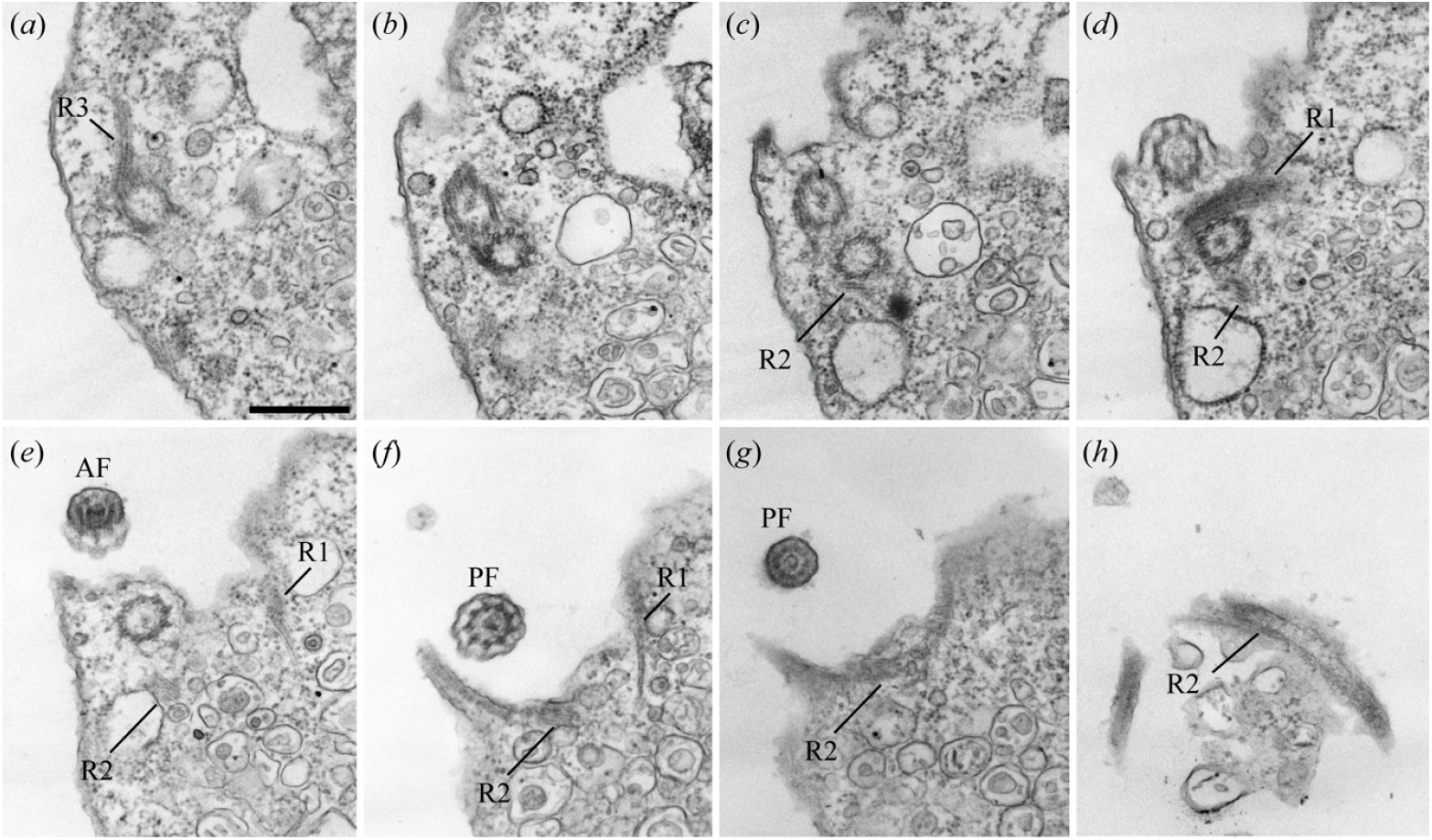
Selected serial transmission electron micrographs of *Glissandra oviformis* n. sp., showing approximately a coronal plane around basal bodies. AF, anterior flagellum; PF, posterior flagellum. Scale bar = 500 nm.

**Figure S3.**
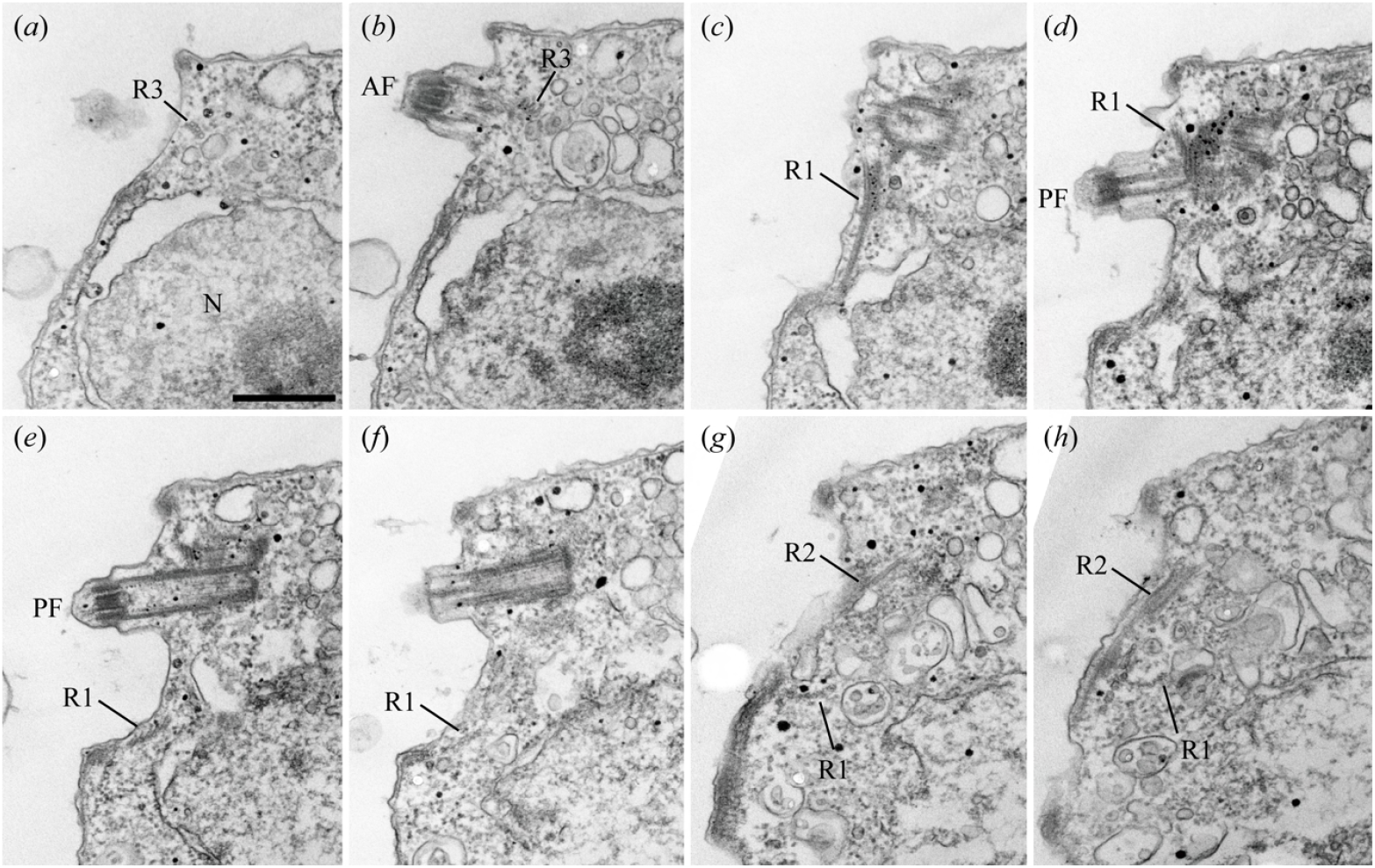
Selected serial transmission electron micrographs of *Glissandra oviformis* n. sp., showing approximately a transverse plane around basal bodies. AF, anterior flagellum; PF, posterior flagellum. Scale bar = 500 nm.

**Figure S4.**
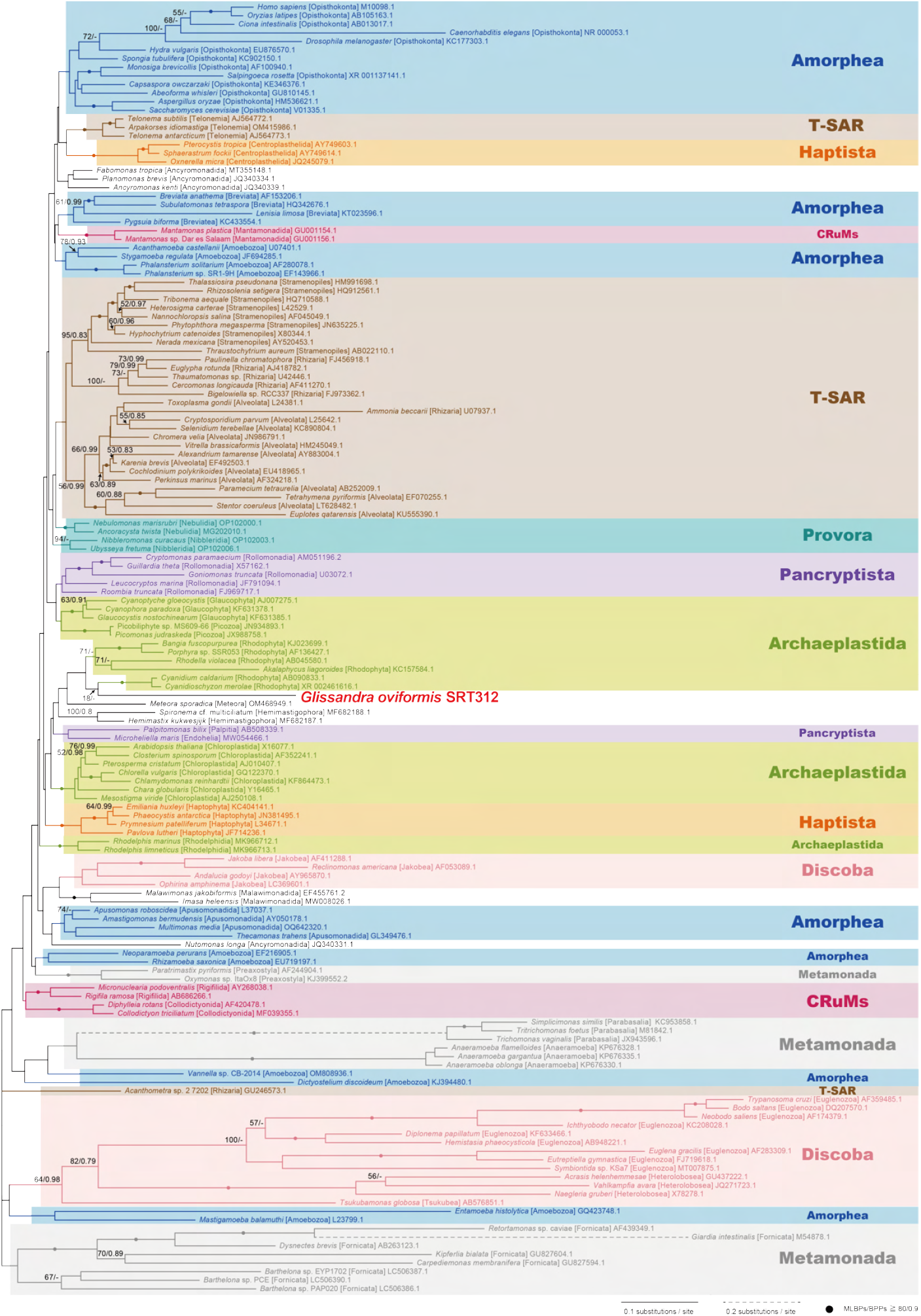
Global eukaryotic phylogeny inferred from small subunit ribosomal DNA (SSU rDNA) sequences. The tree topology was reconstructed using the maximum-likelihood (ML) method. For the support values for bipartitions, ML bootstrap support values (MLBPs) and Bayesian posterior probabilities (BPPs) were mapped onto the ML tree. Nodes marked with dots are supported by MLBPs 80% and BPPs 0.9. MLBPs < 50% and BPPs < 0.8 are not shown unless they are significant to discuss the position of *Glissandra oviformis*.

**Figure S5.**
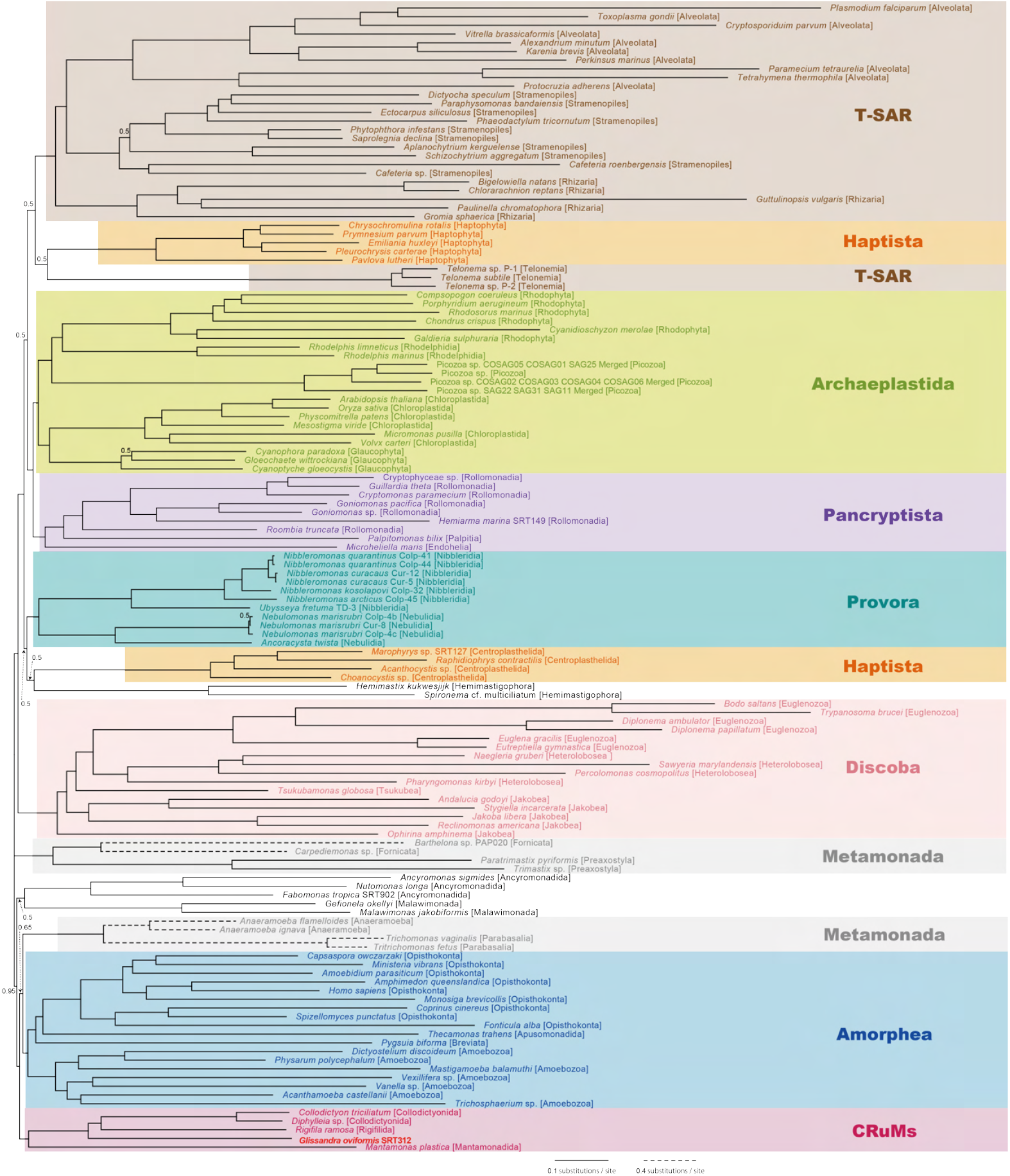
Bayesian phylogenetic inferred from a 340-protein alignment. For each node, Bayesian posterior probabilities (BPP) not equal to 1.0 were presented.

